# New records of signature spiders (Araneidae: *Argiope* spp.) from India with the resurrection of *A. undulata* Thorell, 1887

**DOI:** 10.64898/2026.02.06.704477

**Authors:** Alexander M. Kerr, Samuele Papeschi

## Abstract

We present new distributional records of *Argiope* spiders in India, based on more than 10,000 digital images of the genus from the region curated by iNaturalist (www.inaturalist.org). Notable range expansions to India are documented for three species: *A. chloreides* Chrysanthus, 1961, *A. mangal* Koh, 1991, and *A. sector* (Forsskål, 1776). Second, previously unrecorded field characters, updated distributional data, and a re-examination of published descriptions of type material, support the resurrection of *A. undulata* Thorell, 1887 as a valid species, long treated as a synonym of *A. pulchella* Thorell, 1881. Finally, we report the first *in situ* photographic records of live specimens of the rarely documented *A. caesarea* Thorell, 1897 and *A. macrochoera* Thorell, 1891. These varied findings for a small and conspicuous taxon highlight the value of online community-science platforms for documenting the arachnofauna of a biodiverse region, as well as illustrate the need for continued taxonomic review, even within well-known genera.

## Introduction

The genus *Argiope* Audouin, 1826 (Araneidae: Argiopinae) comprises a globally distributed group of 89 nominal taxa (86 species and three subspecies; see WSC 2025). These large, visually striking orb-weaving spiders inhabiting gardens, roadsides, and native habitats are known for their distinctive and colourful abdominal patterns that make many species readily identifiable in the field. Most, if not all, further stand out by flagging their webs with conspicuous silken decorations of zigzagging crosses and spirals called stabilimenta whose possible function(s) still incite debate (Eberhard 2020; Walter 2023). The genus is considered of ecological and economic importance, as well. Few other spiders possess the broad, strong webs needed to capture the largest of flying insect prey, grasshoppers, beetles, wasps, and moths (Murakami 1985). This and their abundance in agricultural ecotones also make them of interest for their role as biological control agents (Rajeswari and Savitha 2015).

In India, nine species of *Argiope* have been reported to date: *A. aemula, A. anasuja, A. caesarea, A. catenulata, A. lobata, A. macrochaera, A. minuta, A. pulchella*, and *A. trifasciata*. Most of these were first recorded in taxonomic monographs (Pocock 1900; Sinha 1952; Tikader and Biswas 1981; Levi 1983), while regional surveys continue adding state-level records (e.g., Chandra et al. 2021; Sivaperuman and Dash 2022). Like most authorities (e.g., WSC 2025; but see Singh and Singh 2021), we also regard as doubtful the isolated and unprovenanced accounts of the otherwise European *A. bruennichi* (see Leardi in Airaghi 1902), Philippine endemic *A. luzona* (Sankari and Thiyagesan 2010), Sri Lankan *A. taprobanica* (Strand 1907), and Southeast Asian *A. versicolor* (Leardi in Airaghi 1902).

Here, we report four additions to the *Argiope* fauna of India, a 44% increase raising the number to 13 species, and provide the first colour images of live and *in situ* specimens of two rare, previously known taxa. Three of the new records are based on geographic extensions of range and a fourth species is raised from synonymy. Much of this work was enabled by the observations of local “citizen scientists” curated as images by the online database iNaturalist.

## Methods

New distributional records were sourced from digital images submitted by the public to iNaturalist (www.inaturalist.org), an online platform for biodiversity information. We initially retrieved the unique identifiers of all images georeferenced to India by the contributor and assigned by at least one patron to the genus *Argiope* on the iNaturalist API on 8 September 2025 05:30 UTC using JavaScript utilities at http://jumear.github.io/stirfry/. Each observation was manually vetted for species identity based on morphological features found in the taxonomic literature, primarily Levi (1983) and Jäger (2012). For species not reliably distinguishable by habitus alone, we additionally sought images providing a ventral view of adult females in which the epigynum was also in an unobstructed and focussed view allowing comparison with descriptions and illustrations of type material. The pattern on the sternum is also useful (Levi 1983) and was considered when visible. Body length is given as total body length (TBL) of adult females, usually maximum length and taken from the original descriptions or, when also based on type, Levi (1983). In the tallies of *A. undulata* and the leg-colour morphs of *A. pulchella* sensu Thorell, we considered only observations of adult females that displayed at least a dorsal view.

## Results and Discussion

### New range extensions to India

Recent observations curated by iNaturalist have revealed three previously undocumented species of *Argiope* within India. These records extend the known distributions of two Southeast Asian taxa westward and, in one case, a North African and Middle Eastern species further east. In all cases, they represent the first verified national records.

The first, *A. chloreides* Chrysanthus, 1961, was previously known only as far west as Malaysia (Tan et al. 2019) and, like conspecifics elsewhere, appears to prefer well-vegetated habitats, building its web within foliage—often directly atop leaves—and incorporating a disc-shaped stabilimentum (Fig. 1a). The small spider (TBL to 9 mm) is unmistakable in the field (Tan et al. 2019) as the only small green *Argiope* with a banded abdomen (Fig 1b). The 24 independent observations from India indicate its occasional presence in the states of Assam, Arunachal Pradesh, Karnataka, Kerala, Maharashtra, and West Bengal.

**Fig. 1.**
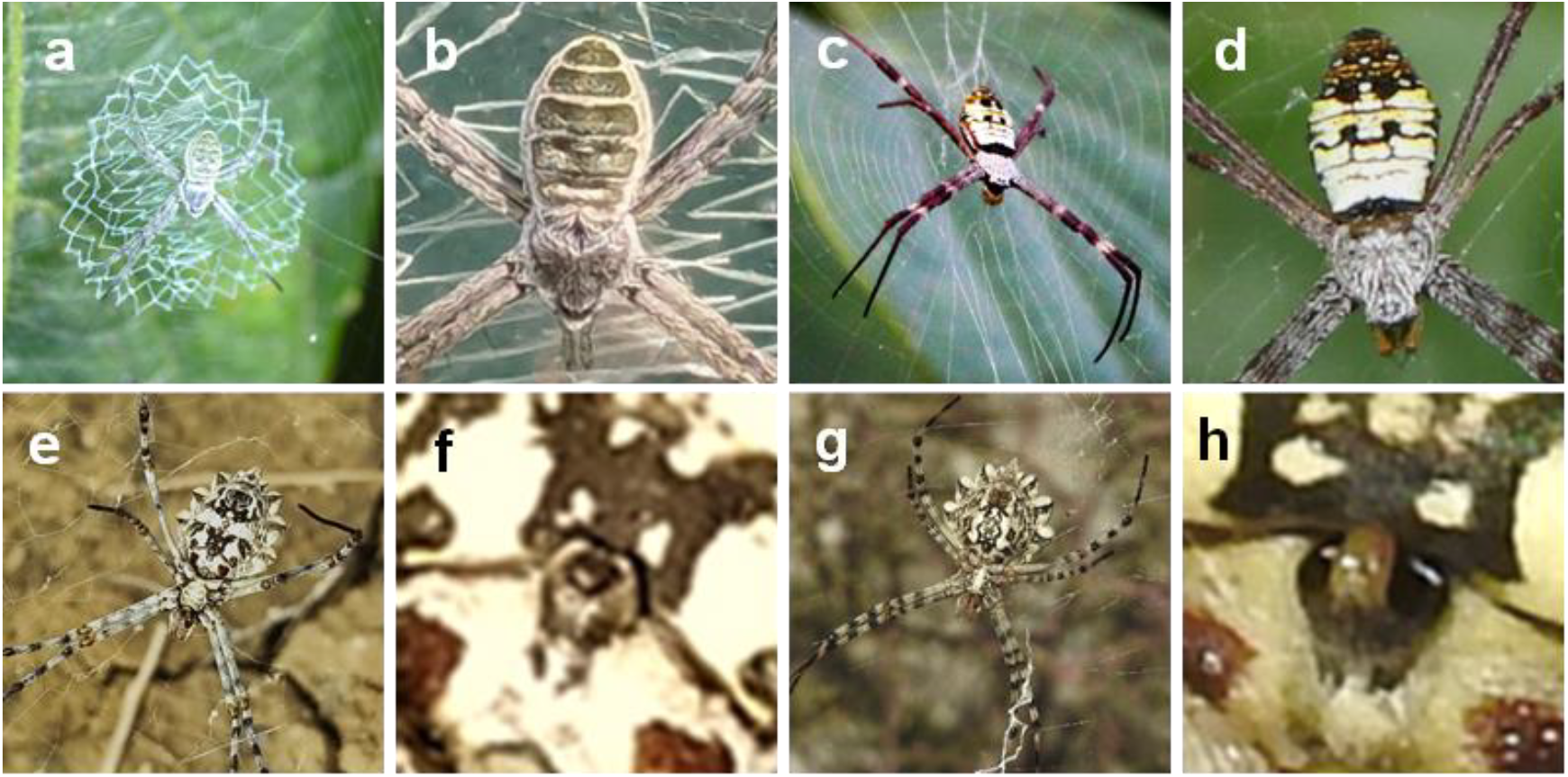
New records of *Argiope* from India. **a)** web built above leaf by *A. chloreides*, Belgaum, Karnataka (261218623), and **b)** abdominal pattern, Changlang, Arunachal Pradesh (201460934). **c)** *A. mangal* in web, Bally, West Bengal (112350777), and **d)** abdominal pattern, North 24 Paragnas, West Bengal (114273140). **e)** *A. sector*, Junagadh, Gujarat (245251202), and **f)** epigynum; **g)** *A. lobata*, Spain (248741740), and **h)** epigynum. Image numbers from iNaturalist (www.inaturalist.org); photographers: **a)** Raghavendra H.K., **b)** Rohan Menzies, **c)** Abhik Rong, **d)** Anubhav Agarwal, **e–f)** Jignesh Rathod, **g–h)** Víctor Javier Marugán; licencing: **a, c–d, g–h)** CC-BY-NC, **b)** CC-BY, **e–f)** all rights reserved with permission

The second species, *A. mangal* Koh, 1991, is somewhat larger (to 12 mm TBL) and was previously reported only from its type locality in Singapore, where the diagnostic abdominal pattern is yellow or white (Koh 1991); to date only white has been seen in the few Indian specimens (Figs. 1c–d). As in Singapore, the species appears to be restricted to coastal mangrove habitats. We verify its presence via two sightings in India from West Bengal.

The third species, *A. sector* (Forsskål, 1776), previously known only as far east as Iran (Mirshamsi et al. 2015), is reported here for the first time from India, based on two records from Gujarat (Fig. 1e–f). Unlike the previous two species, *A. sector* lacks diagnostic gross morphological characters useful for field identification that have been corroborated by genitalic features and nucleotide sequence data (Tan et al. 2019). Rather, *A. sector* is indistinguishable by overall appearance alone from *A. lobata*, another large (to 24 mm TBL) Indian congener with a pronounced multilobate abdomen. However, *A. sector* is the only *Argiope* whose epigynum (Fig. 1f) possesses a broad, coarse roof hiding the atrial openings that is topped by a large, often darkened shiny knob (Bjørn 1997). This contrasts with *A. lobata* (Fig. 1g) whose epigynal roof lacks a knob and is smooth, shiny, and narrow so that the black, round atrial openings are plainly visible (Fig. 1h). The two records of *A. sector* from India suggest that it is either a genuinely rare addition to the country’s arachnofauna, or that perhaps it has been overlooked in photographic records and misidentified as *A. lobata*. However, the latter species also appears to be uncommon; of the only 24 observations from India on iNaturalist, the five adult females with clear views of the epigynum indicate none approach that of *A. sector*.

### Resurrection of *Argiope undulata* Thorell, 1887

Our review of Thorell’s original description and the subsequent literature illustrating type material support the reinstatement of *A. undulata* as a valid species. Although synonymised without explanation under *A. pulchella* Thorell, 1881 by Levi (1983), *A. undulata* can be reliably distinguished even in the field by multiple morphological characters. We therefore formally resurrect this species and present evidence clarifying its diagnostic features and known distribution.

Thorell (1887) and Levi (1983) both describe *A. undulata* as a large *Argiope*, the adult female TBL to 29 mm. The authors both relate the syntypes’ eponymous undulating lateral and posterior margins of the abdomen, dorsally three broad, light-coloured, often yellowish in life, and widely spaced transverse bands alternating upon a grey to olive-brown ground often with scattered light speckles (Fig. 2a–g). By contrast, *A. pulchella* sensu stricto (s. s., in the original sense meant by Thorell) is smaller (TBL to 20 mm), has a more regular pentagonal abdomen lacking an undulating margin unless post-gravid, and exhibits a more complex dorsal abdominal pattern involving more colours and larger white spots, the ground colour usually reddish-brown to black.

**Fig. 2.**
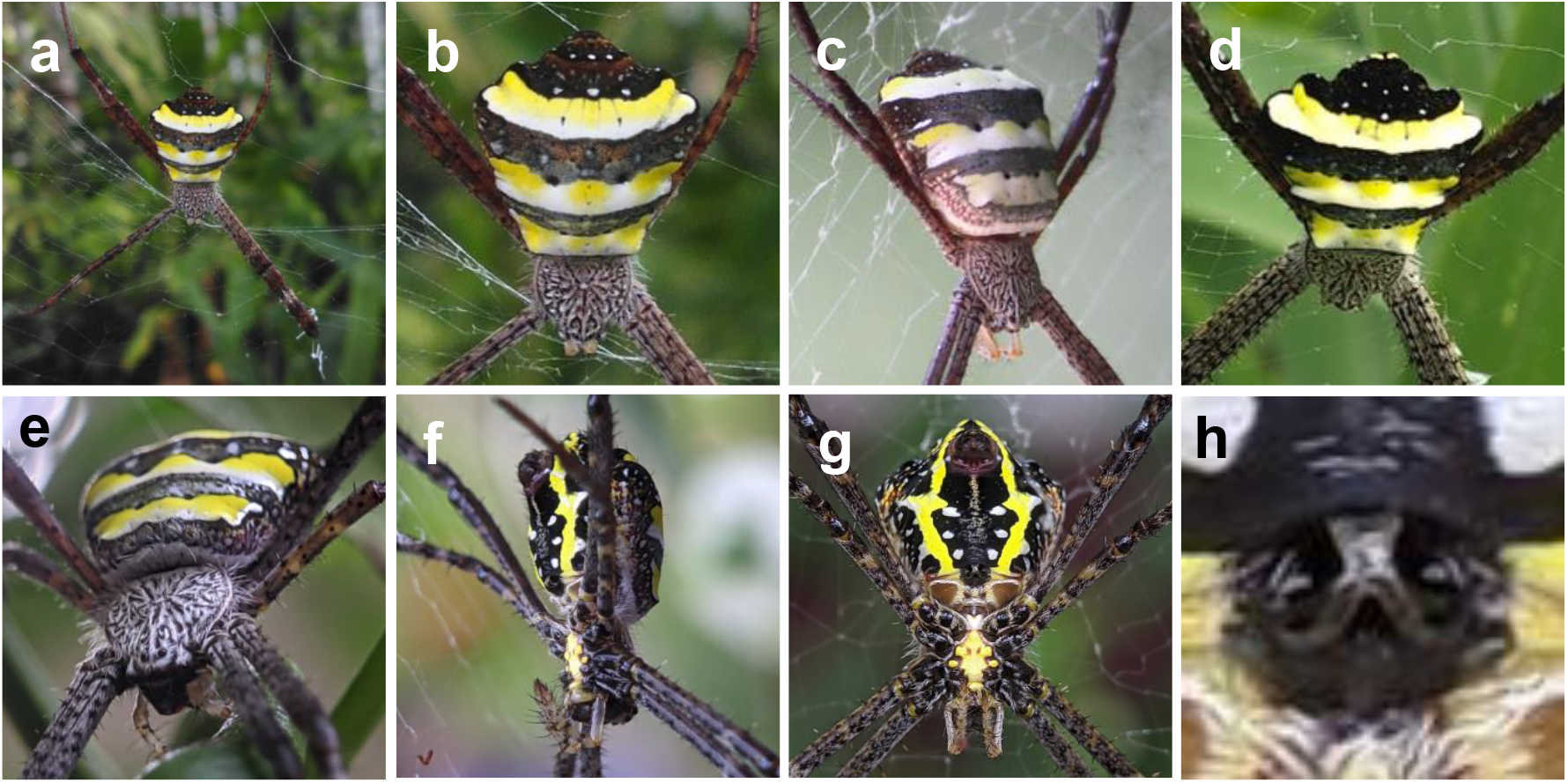
*Argiope undulata* Thorell, 1887. **a)** Habit and abdominal patterns, **b)** Malappurram, Kerala (181043145), **c)** Chennai (185875459) and **d)** Ciombatore (257224487), Tamil Nadu; **e–g)** multiple views of body and **h)** epigynum, Kannur, Kerala (177339177). Image numbers from iNaturalist (www.inaturalist.org); photographers: **a–b)** athifkenz **c)** chinmaya30, **d)** milestoswim, **e–h)** Vaishnav K. V.; licencing: **a–c)** CC-BY-NC, **d)** CC-BY, **e–h)** CC-BY-SA

Further, as illustrated by Levi (1983: Figs. 238–243) in a side-by-side comparison of type and discussion by Pocock (1900) of non-type, the epigyna of these species bear little resemblance to one another. The epigynum of *A. undulata* (Fig. 2h) has a more sloping roof indented medially and partially covers the atrial openings and septum in ventral view, with somewhat thickened rounded posterior rims divided by a distinct septum that widens posteriorly to form the posterior wall or plate. By contrast, the epigynum of *A. pulchella* s. s. has a dome-like roof nearly to fully covering the openings and septum, with a uniformly thicker and rounded posterior rim and a shorter, narrower posterior plate (see Figs. 238–240 in Levi 1983; Figs. 56, 58, 61, 63, 65, 70 in Jäger and Praxaysombath 2009).

Additionally, specimens matching *A. undulata* invariably lack a trait private to *A. pulchella* s. s. across the latter’s entire range (Fig. 3): strongly banded legs in 20.9% of iNaturalist images of adult females of this species from India (*n* = 1048)—first remarked upon by Pocock (1900) and observed in some other *Argiope* species (e.g., Friedman 2025). Finally, unlike *A. pulchella* s. s., which occurs widely across India excepting the driest western region, *A. undulata* appears to be primarily an endemic of the Eastern and Western Ghats ranges (Fig. 3), occurring northward in similar environments to at least Myanmar, its type locality. Hence, the species looks to be restricted in India to the states of Andra Pradesh, Goa, Kerala, Karnataka, Maharashtra, Northeast India, Odisha, Telangana, Tamil Nadu, and West Bengal.

**Fig. 3.**
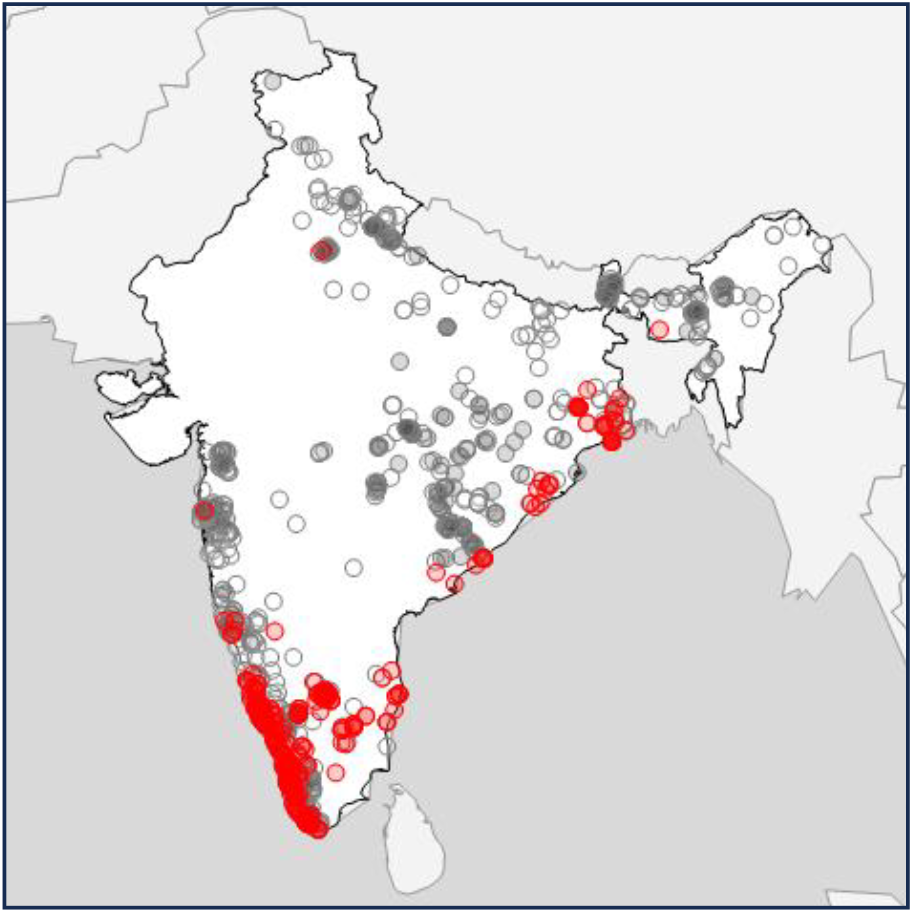
The distributions in India of *Argiope undulata* (red circles; *n* = 311) and *A. pulchella* sensu stricto with banded (grey circles; *n* = 219) and unbanded legs (open circles; *n* = 829). Specimen images and locality data from iNaturalist (www.inaturalist.org).

In sum, we therefore regard *A. undulata* as a valid species and remain uncertain why Levi (1983) synonymised the nomen without comment. In fact, at least one early author (Strand 1907) likely regarded the species as closer to the presumed Sri Lankan endemic *A. taprobanica*, an issue we at present lack data to address.

### Photo vouchers of rare species

Among the previously known Indian *Argiope*, at least two species have notably restricted distributions within the country and are seldom reported (Singh and Singh 2021): *A. caesarea* in Northeast India and *A. macrochoera* in Andaman and Nicobar. Here, we present image-based records of both species live and *in situ*, the first to our knowledge.

*A. caesarea* Thorell, 1897 (21–26 mm TBL) was originally described from Myanmar, and later reported from Northeast India (Sinha 1952; Tikader 1970, 1982) and West Bengal (Chandra et al. 2021). Photographs from India (Figs. 4a–d) depict a large, distinctive form consistent with Thorell, as well as Levi’s (1983) and Xiaoqui et al.’s (2024) redescriptions: yellow palps, black legs, and a sub-pentagonal abdomen with smooth lateral and posterior margins, dorsally dark with three broad transverse pale yellow bands. The anterior two pale bands are separated by a narrow yellowish brown stripe, while the third band is recurved and set off from the second by a wider field that can range widely in colour among individuals from black (Fig. 4a) to yellowish brown (Fig. 4b). The sternum is black with a narrow medial yellowish streak (Fig. 4c). The epigynum is likewise diagnostic: unusually broad and blunt, the wide posterior plate and roof nearly covering the atria (Fig. 4d).

**Fig. 4.**
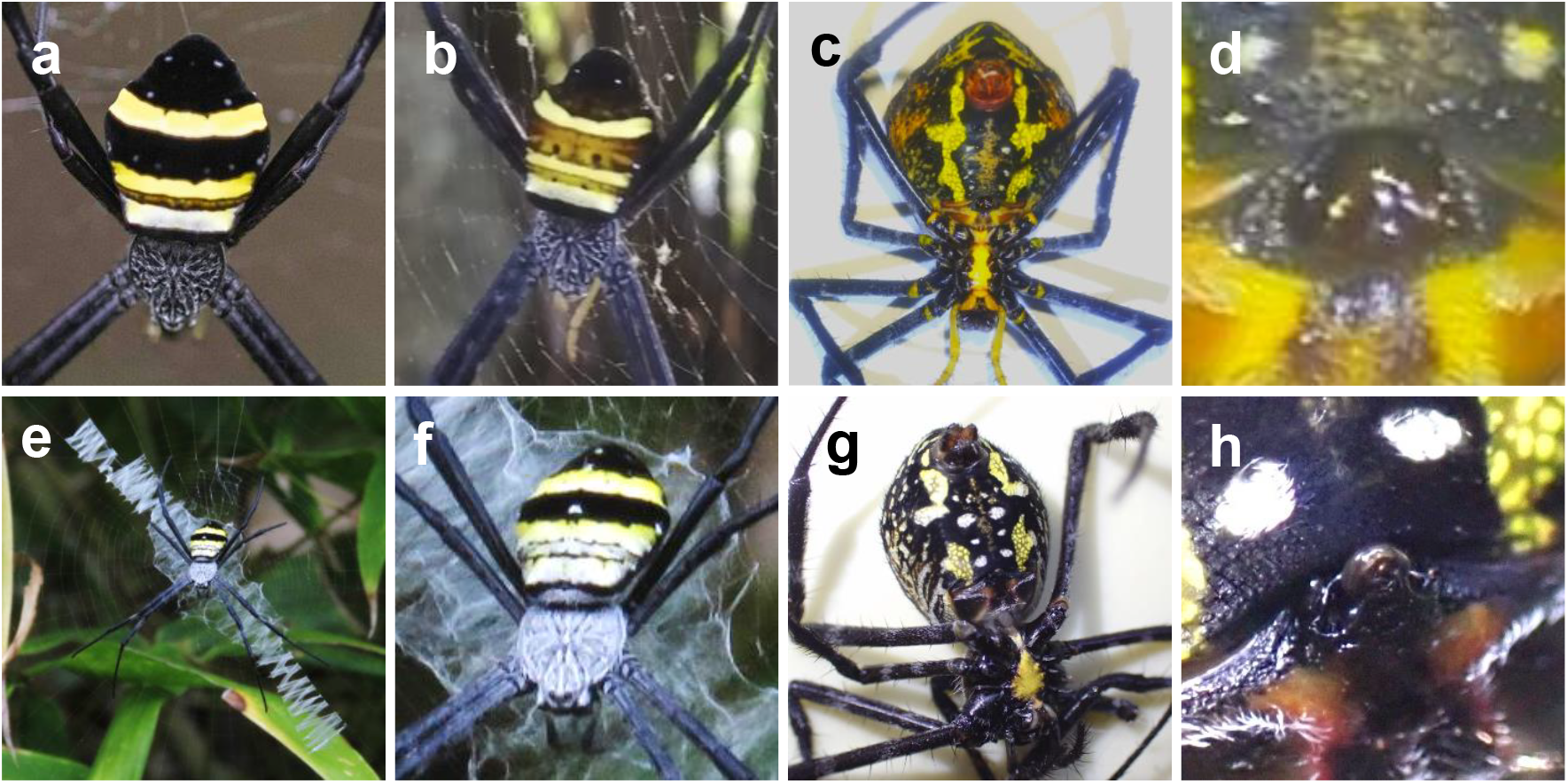
Photo vouchers of rare Indian *Argiope* spp.: *A. caesarea* Thorell, 1897, showing variation in abdominal colour from **a)** Darjeeling, West Bengal (253986786) and **b)** East Sikkim (244263692), and the **c)** ventrum and **d)** epigynum, Darjeeling (304468940); *A. macrochoera* Thorell, 1891, Great Nicobar, Andaman and Nicobar, showing **e)** habit, **f)** dorsum (104906322), **g)** ventrum, and **h)** epigynum (327060942). Image numbers from iNaturalist (www.inaturalist.org); photographers: **a)** amolmm, **b)** Kiron Dey, **c–d)** rodha_rai-3, **e–f)** zankhna, **g–h)** Sudharsan E.; licencing: **a, c–h)** CC-BY-NC, **b)** CC-BY-NC-ND

The second recently photographed species, *A. macrochoera* Thorell, 1891 (at least 14 mm TBL), is an endemic of Andaman and Nicobar (Sivaperuman and Dash 2022). Observations from Great Nicobar Island (Figs. 4e–h) depict a species matching Thorell’s and Levi’s descriptions, having black, rather spiny legs and a pentagonal-ovate abdomen, thus longer than broad, dorsally black with very few small spots, overlain by an anterior light-coloured trapezoid divided by one or two thin, transverse dark lines, and more posteriorly a broad central light transverse band (Figs. 4e–f). The sternum is completely yellow, the tibiae of leg pair IV are densely setate, and the ventrum, unlike for all other *Argiope*, displays two to three pairs of short paraxial streaks, one with short transverse extensions pointing laterally (Fig. 4g). The epigynum is also unique, broad with the rim jutting posteriorly beyond the septum as an elevated, rounded scape (Fig. 4h), reminiscent of some non-argiopine araneids, such as *Neoscona* (see, e.g., Zamani et al. 2020).

## Conclusions

This small trove of new and revised national records underscores the value of open-access biodiversity platforms, which have accelerated the accumulation of natural-history data, particularly in nations with a wealth of biodiversity, such as India. Still, we were struck that even within a well-known genus of moderate global diversity like *Argiope*, a nationwide review could yield such a relatively large haul of newly documented species. Remaining questions include ascertaining the relationship of *A. undulata* to *A. taprobanica* and resolving the long-recognised uncertainty around the concept of *A. pulchella*. Community involvement will continue to be a valuable addition to these efforts, as well as in revealing further and perhaps even new species of *Argiope* in India and elsewhere.

## Acknowledgements

We thank the iNaturalist community of developers and contributors for transforming the role of natural history in science, the Biodiversity Heritage Library, World Spider Catalog, and the Natural History Museum of Bern for access to the early literature, iNaturalist observers Anja Junghanns (University of Greifswald), einsum (unaffiliated), Naufal Urfi Dhiya’ulhaq (SpeciesObscura.org and University of Göttingen) for discussion and sorting thousands of image identifications of *Argiope*, as well as Sudharsan E. (Zoological Survey of India) for fieldwork on *A. macrochoera*. This paper is a contribution of The Marine Laboratory, University of Guam.

